# Zika virus replicates in skeletal muscle contributing to peripheral viral amplification prior to reach neural tissue

**DOI:** 10.1101/2020.03.26.010389

**Authors:** Daniel Gavino-Leopoldino, Camila Menezes Figueiredo, Letícia Gonçalves Barcellos, Mariana Oliveira Lopes da Silva, Suzana Maria Bernardino Araújo, Rômulo Leão da Silva Neris, Laryssa Daniele Miranda, Leandro Ladislau, Claudia Farias Benjamim, Andrea Thompson Da Poain, Julia Rosauro Clarke, Claudia Pinto Figueiredo, Iranaia Assunção-Miranda

## Abstract

Zika virus (ZIKV) infections are still a worldwide concern due to the severity of neurological outcomes. ZIKV neurotropism is well characterized, but peripheral tissue could be sites of viral amplification, contributing to endothelial-barrier crossing and access to peripheral nerves. During acute and late phases of infection, ZIKV can be detected in several body fluids, eyes, testis and vagina. However, the importance of initial replication sites for the establishment of infection and viral spread remain unknown. Here we demonstrated that ZIKV replicates primarily in human muscle precursor cells, resulting in cell death and inhibition of myogenesis. ZIKV also replicates in fetal muscle after maternal transmission and in infected neonate mice, inducing lesions and inflammation. Muscle was an important site of viral amplification, sustaining higher peripheral viral loads than liver and spleen. In addition, ZIKV showed rapid and sustained replication kinetics in muscle even before replication in the neural tissues, persisting until 16 days post infection. Our results highlight the importance of muscle in ZIKV pathogenesis as a peripheral site of viral amplification which may contribute to ZIKV reaching neural structures.

**Author Summary:** Zika Virus (ZIKV) neurotropism and its deleterious effects on central nervous system have been well characterized. But, investigations of the initial replication sites for the establishment of infection and viral spread to neural tissues remain under explored. Here we demonstrated that ZIKV replicates primarily in human skeletal muscle precursor cells, resulting in cell death and disrupted myogenesis. ZIKV also replicates in muscle of fetus and neonate mice inducing muscle damage and inflammation. Muscle replication occurs before amplification in peripheral nerves and brain, contributing to the increase of peripheral ZIKV load and dissemination. In addition, ZIKV RNA still been detected in skeletal muscle at late stages of infection. Overall, our findings showed that skeletal muscle is involved in ZIKV pathogenesis, contributing to a broader understanding of ZIKV infection. Thus, opens new aspects in the investigation of the long-term consequence of early infection.

## Introduction

Zika virus (ZIKV) is an arbovirus from the *Flaviviridae* family transmitted mainly by *Aedes* mosquitoes. ZIKV disseminated rapidly across the Americas in the last years (1) and although historically this infection caused a self-limited mild febrile condition similar to that induced by other arboviruses, reports of persistent and severe neurological damage started to appear. It is now known that vertical transmission of ZIKV can cause fetal death and congenital defects, such as microcephaly (2) while neuroinflammatory conditions such as Guillain-Barré Syndrome (GBS) and encephalitis have been reported in adult patients (3-6).

Several studies have described the molecular mechanisms of the pathogenesis of ZIKV infection, mainly neuroinvasion and the damage on the central nervous system (CNS). It has been demonstrated that ZIKV replicates in human neural progenitor cells and brain organoids, resulting in cell death and cell cycle arrest (7, 8). Both intra-uterine and neonatal ZIKV exposure in mice was shown to compromise neurogenesis and cause severe necrosis in different brain regions, along with persistent viral replication and widespread neuroinflammation (9, 10). Moreover, it was shown that activation of microglial cells and astrocytes associated to viral replication in the brain affects cognitive function and the differentiation of glial progenitors cells impairing brain development (11, 12).

Under physiological conditions, the blood brain barrier (BBB) protects the brain from the effects of peripheral infections. However, in some circumstances the integrity of this endothelial barrier can be broken allowing pathogenic agents to reach the brain. It has been hypothesized that viral replication in peripheral tissues could contribute to amplification of viral loads early during ZIKV infection and before the virus invades the CNS (13). ZIKV has been shown to replicate and cross the endothelial barrier without significantly affect its permeability (14, 15). Additionally, it was demonstrated that ZIKV replicates in human peripheral neurons *in vitro* (16). Thus, replication in motors and sensory tissues, in addition to contributes to viral load, could be a route used by ZIKV to access the peripheral nerves and reaches by retrograde axonal-transport some immuno privileged sites such as the brain (17). This is a mechanism known to be involved in infections by other neurotropic viruses including the member of Flaviviridae family, West Nile virus (18-20). Despite the evidence that ZIKV can be detected both during acute and late phases of infection in several body fluids, eyes, testis and vagina (21). The importance of initial replication sites for the establishment of infection and viral spread remain unknown.

Here we investigated the ability of ZIKV to establish a productive replication in the skeletal muscle. We found that ZIKV replicates in human muscle precursor cells, impairing differentiation and causing cell death. Using mouse models, we showed that ZIKV induce necrotic lesions and inflammation in the muscle of fetuses and newborns by establishing rapid and sustained replication kinetics. Interestingly, ZIKV replication in the muscle temporally precedes the detection of viral RNA in the brain. Taken together, our results indicate that the skeletal muscle is an early site of viral amplification that may contribute to ZIKV reaches neural tissues.

## Results

### ZIKV replicates in human skeletal muscle progenitor cells, causing cell death

The main extra-neuronal sites of ZIKV replication and the role of peripheral viral replication in the establishment of infection are still poorly understood. In order to evaluate whether muscle is a potential site of ZIKV replication, we infected primary human skeletal muscle myoblast cells (HSMM), both in an undifferentiated stage (myoblast culture) and after cell fusion and differentiation into muscle fibers (myotubes). Temporal quantification of infectious particles in culture media (**Fig 1A**) revealed a rapid increase in ZIKV infectious particles in myoblast cultures, which lasted for at least 48 hours post-infection (hpi). No increase in viral particles from myotubes in culture was observed until 16 hpi, but titers were comparable to those seen in myoblasts at 36 hpi. In agreement with the replication curves, we positive immunostaining against ZIKV E protein (**Figs 1B and 1C**) and ZIKV non-structural 2B protein (NS2B) (**Figs 1D and 1E**) in both myoblasts and myotubes in culture, when analyzed 36 hpi. No staining for either viral protein was observed in mock myoblast (**S1A Fig**) and myotubes (**S1B Fig**) cultures. Quantification of 4G2-positive cells showed a significantly higher number of ZIKV-positive cells in myoblast cultures when compared to myotube cultures at 36 and 48 hpi (**Fig 1H**).

**Figure 1.**
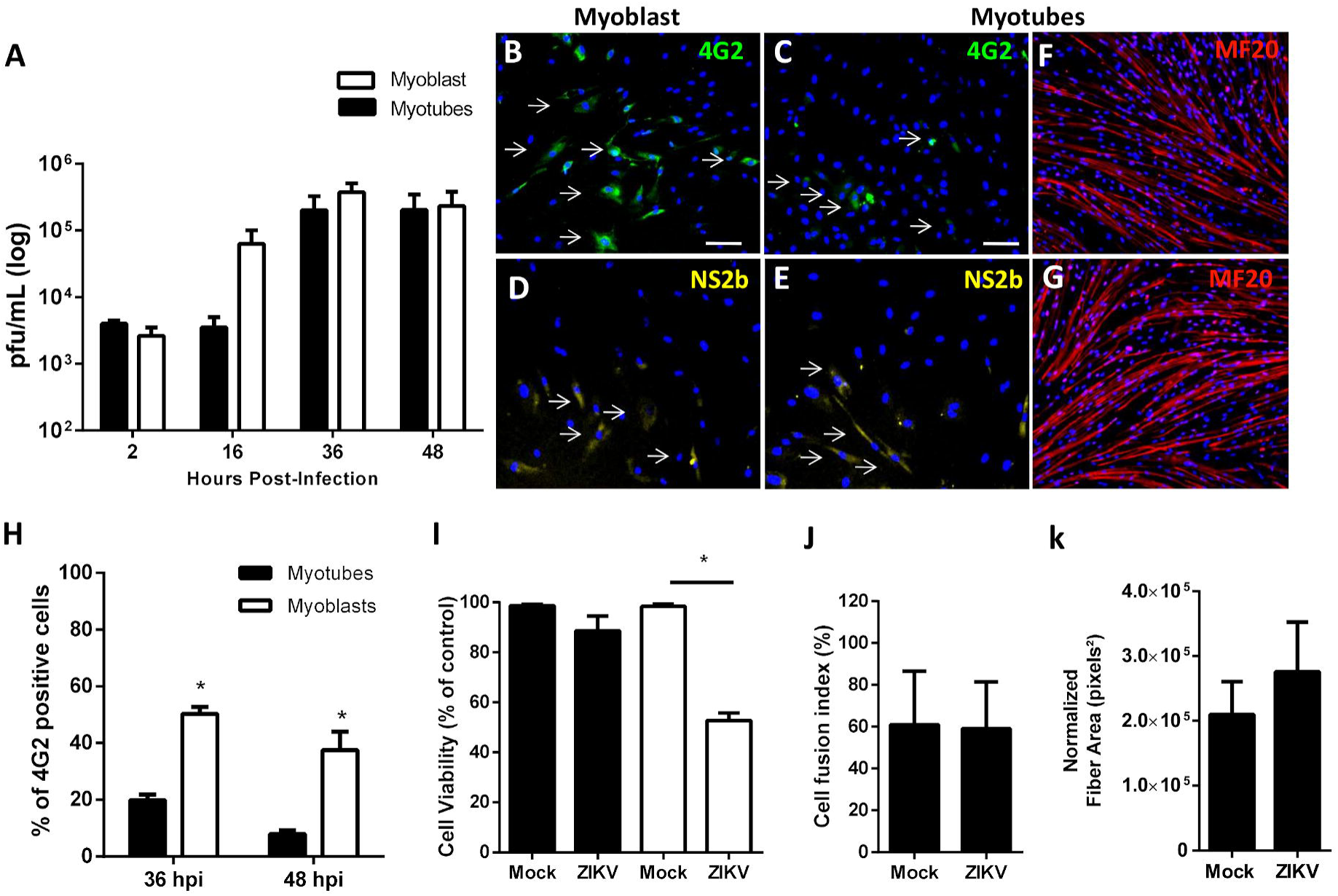
ZIKV replicates and promotes death of human skeletal muscle progenitor cells. Human primary skeletal myoblast and differentiated myotubes culture were infected with ZIKV with a MOI of 5 and temporally assessed. (**A**) ZIKV released at cultures supernatant at different times post-infection was quantified by plaque assay (n=3-5 each point); (**B-G**) Immunofluorescence analysis for detection of ZIKV positive myoblast (**B and D)** and myotubes (**C and E**) 36 hours post-infection using anti-flavivirus E protein 4G2 and anti-NS2b of ZIKV. White arrows indicate some positive cells; (**F-G**) detection of fibers at Mock (**F**) and ZIKV (**G**) infected myotube culture using anti-myosin heavy chain (MF20) 48 hours post infection; (**H**) Quantitative analysis of 4G2 positive cells in % of total (Nuclear staining with DAPI) was performed using at least 10 fields of 2 independent experiments using ImageJ software; (**I**) myoblast (white bars) and myotubes (black bars) viability was determined by MTT reduction 48 hours post ZIKV infection related to Mock of 3 independent experiments in triplicate; (**J-k**) Quantification of MF20 positive was used to determine cells fusion index (**J**) and fiber area (**k**) related to total of cells (nuclear staining with DAPI) at differentiated culture 48 hours post ZIKV infection and mock using at least 10 fields of 2 independent experiments using ImageJ software. Data was analyzed using ANOVA (**A and H**) and Mann-Whitney non-parametric tests (I). *p□0.05

Next, we evaluated whether ZIKV replication affects the viability of skeletal muscle cells. When analyzed 48 hpi, we observed that while infection promoted reduced myoblast viability to ∼50%, viability of myotubes in cultures exposed to ZIKV was not affected (**Fig 1I**). Furthermore, consistent with these observations, myosin heavy chain (MF20) immunolabeling in mock- and ZIKV-treated myotube cultures showed that ZIKV infection did not alter fiber structure when compared to cells treated with mock (**Figs 1F and 1G**). In addition, quantification of MF20 images showed that the infection did not altered the number of fibers nor the fiber area (**Figs 1J and 1L**), suggesting that ZIKV infection does not interfere with the integrity of the already formed myofiber structures. Taken together, these data suggested that ZIKV is capable of replicating in myogenic cells causing significant cell damage.

### ZIKV infection inhibits myogenesis

Considering our results showing that ZIKV replicates and causes death of muscle progenitor cells, we investigated whether ZIKV interferes with myogenesis by altering the number and integrity of the newly formed fibers. To this end, we subjected human myoblast cultures to fusion and differentiation-inducing medium and infected these cells at an early stage of differentiation (1-day post-differentiation stimuli). Fiber formation was evaluated by MF20 immunolabeling after infection (day 1) and at 5-day post-differentiation stimuli, that cultures reached late stage of differentiation (day 5), and showed a reduction of MF20 positive cells in ZIKV infected culture at day 5 comparing to Mock (**Fig 2A**). Quantitative analysis indicated that ZIKV infection promoted a decrease in the average fiber-area and in the amount of differentiated fibers, 5 days after differentiation induction (**Figs 2B and 2C**). We also observed a rapid increase in infectious viral particles in culture medium of ZIKV-infected myoblasts under differentiation, and ZIKV replication was maintained for at least 96 hpi (**Fig 2D**). We also observed a reduction in MTT metabolization when compared to mock-treated cultures (**Fig 2E**). These data showed that myoblast differentiation and fiber-occupied area were impacted by viral exposure, suggesting that myogenesis is directly affected by ZIKV infection.

**Figure 2.**
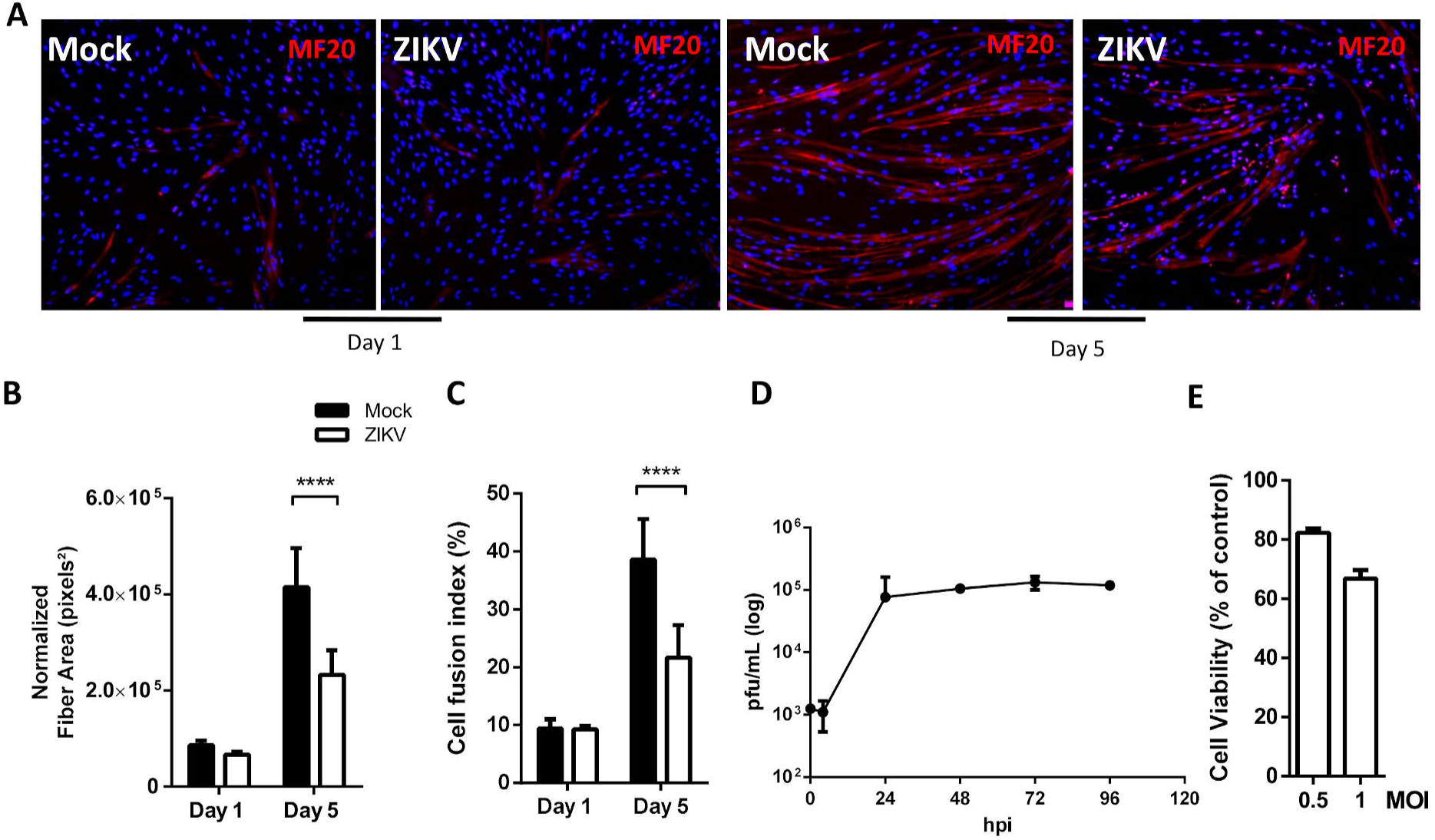
ZIKV infection inhibits myogenesis. Skeletal muscle progenitor cells were subjected to differentiation into myotubes. At day 1 of differentiation were Mock or ZIKV infected with MOI of 1, then was cultured until day 5 of differentiation. (**A**) Cells were fixed at day 1 and 5 of differentiation and formed fibers were detected by immunofluorescence using MF20 (red) and cell nucleus was stained with DAPI (blue). (**B**) Fiber area and (**C**) fusion index (%) of mock (black bars) and ZIKV infected cell (white bars) were obtained by fluorescence images analysis of at least 10 fields of tree independent experiments using ImagJ software. (**D**) ZIKV released at culture supernatant at different times post-infection was quantified by plaque assay (n=3, each point). (**E**) Cell Viability was determined by MTT metabolization 96 h post ZIKV infection with MOI of 0.5 and 1; values are expressed as a % of mock group (control). Statistical analysis was performed by two-way ANOVA. ****p□0.0001

### ZIKV replicates and promotes damage in the skeletal muscle in vivo

Since myoblast fusion is critical not only for the correct development of skeletal muscle but also for the adequate function of the developed tissue (22), we investigated the ability of ZIKV to replicate and induce muscle tissue damage during early stages of development *in vivo*. To this end, we first used a mouse model of ZIKV maternal transmission using type-I interferon receptor deficient (IFNAR-/-) pregnant mice in different gestational stages (**Fig 3A**). ZIKV subcutaneous infection at 10.5 days of pregnancy resulted in 100% fetal loss (data not show), whereas a significant reduction in the number of pups born from each dam was observed when infection was conducted at 12.5 days of gestation (**Fig 3B and S2A Fig**). Fetal death following ZIKV infection was further confirmed by the presence of fetal resorption fragments found in the placenta of infected mice (**S2B Fig**). We also observed difference in the pups, which showed low motility and cases of stillbirths after maternal infection in 12.5 days of gestation (**S2A Fig**). The number pups per offspring was not affected when infection was performed at either 14.5 or 18.5 days of gestation. ZIKV RNA was quantified in pups’ skeletal muscle from left and right hind legs collected bilaterally at post-natal day 0. Although muscular viral titers at birth were higher when infection was performed at earlier stages of pregnancy, viral RNA was detected after infection at all gestational tested (**Fig 3C**). ZIKV RNA was also detected in brains of newborn pups (**Fig 3D**), and viral load was comparable between brain and muscle. Fetal fragments collected from placenta of dams infected at 12.5 days of gestation, however, showed higher viral loads than both brains and muscles of pups analyzed on the day of birth (**Fig S2B**). These data indicate that muscle is a site of ZIKV replication during embryogenesis, sustaining detectable viral loads until the birth of pups.

**Figure 3.**
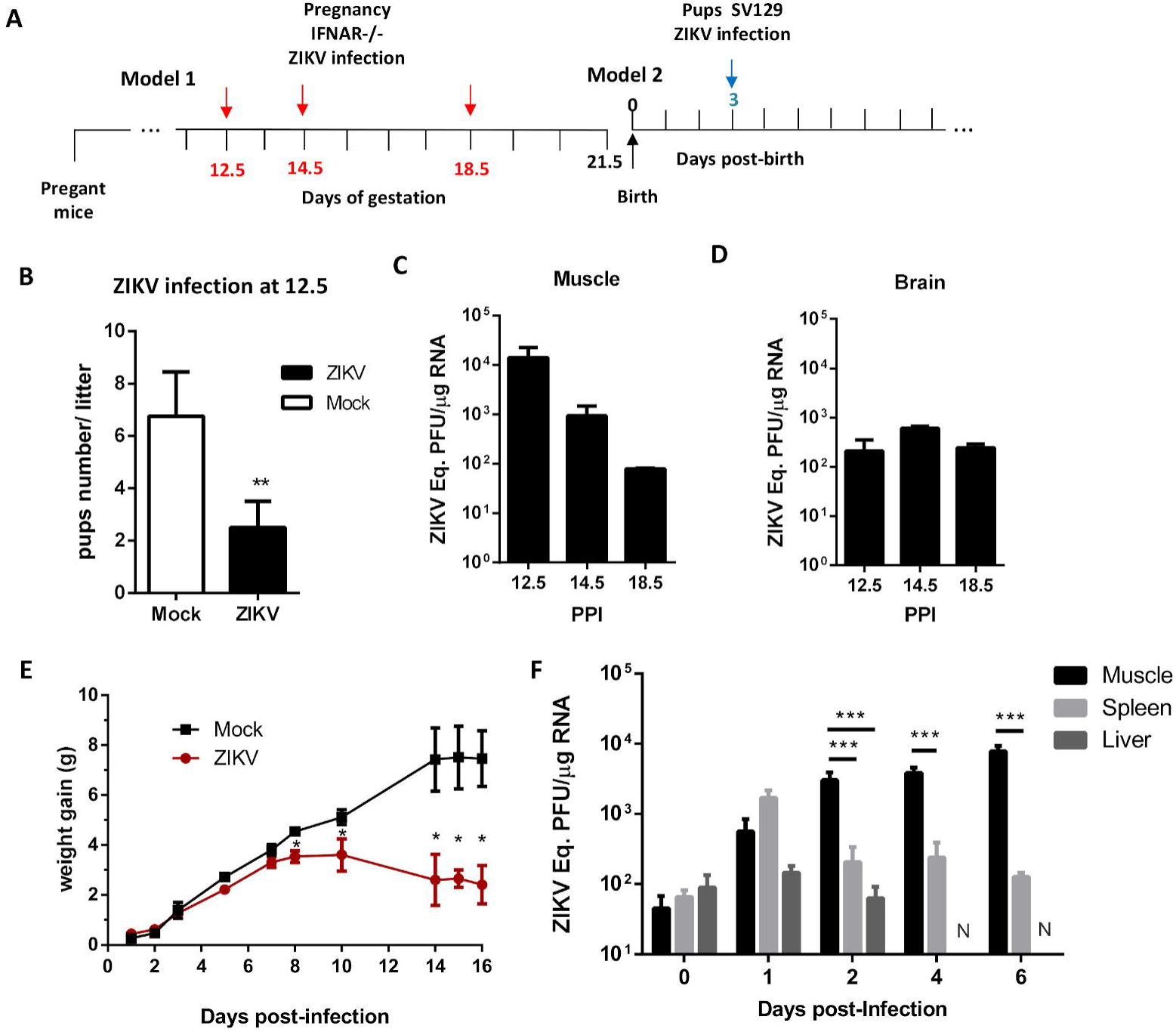
ZIKV replicates at mice muscle after maternal transmission and neonatal infection. (**A**) Schematically representation of the period of ZIKV infection at pregnant SVA129 mice (**model 1 – red arrow**) and pups of SV129 mice (**model 2 – blue arrow**). **Model 1** - SVA129 female was Mock or ZIKV infected (10^5^ pfu) at different pregnancy period (12.5, 14.5, and 18.5, n=3 each) and pups were analyzed at birth; (**B**) Number of pup per litter after infection at 12.5 of pregnancy; (**C and D**) Muscle and brain were collected at birth after ZIKV inoculation at different pregnancy period of infection (**PPI**). ZIKV RNA at tissue was quantified by qPCR in samples of 3 independent experiments. **Model 2 –** 3 days-old WT SV129 mice was mock or ZIKV infected (10^6^ pfu) and temporally accompanied. (**E**) Weight gain after infection (n=8); (**F**) muscle, liver and spleen tissues were collected to quantification of ZIKV RNA load by qPCR at different times post infection (n=4-6, each point). N: indicates non-detectable. Statistical analysis was performed by (**I**) unpaired t test and (**E and F**) two-way ANOVA followed of Tukey’s multiple comparisons test. *p□0.05, **p□0.01 and ***p□0.001.

We also investigated ZIKV replication in 3 days-old wild-type (WT) neonatal mice, infected subcutaneously with ZIKV. We observed that ZIKV-infected mice presented a significant reduction in body weight gain when compared to the mock group (**Fig 3E**) and a high mortality rate starting 12 days after ZIKV inoculation (**S2C Fig**). The quantification of ZIKV RNA in the skeletal muscle form hind leg of mice collected bilaterally showed a rapid increase in the viral titers after inoculation (**Fig 3F**), with the replication peak occurring at 2 days post-infection (dpi), and plateauing at least until 6 dpi. We also observed an increase of ZIKV titers in spleen at 1 dpi, but in this case the titers decreased to a very low level in the subsequent days (**Fig 3F**). In addition, we did not observe a temporal increase in ZIKV titers in the liver, which showed undetectable values after 4 dpi (**Fig 3F**). These observations indicate that, compared to the rodent spleen and liver, the muscle is an important site of ZIKV amplification amongst peripheral tissues in mice, being able to sustain high viral loads for longer periods.

We next investigated whether ZIKV replication in muscle triggers tissue injury and inflammation. Histological analysis demonstrated that ZIKV infection is accompanied by an inflammatory infiltrate in the muscle of newborn pups (P0) after maternal transmission and of neonatal mice 6 dpi (**Figs 4A-4F**). In addition, we observed necrotic areas and alterations on fiber structure. We also found indications of muscle atrophy (**Figs 4B and 4C**). In agreement with the observed morphological alterations, we found increased levels of mRNA of several pro-inflammatory mediators, including TNF, IL-1β, IL-6 and RANTES, in the skeletal muscle of ZIKV-infected mice compared to mock-injected mice at 6 dpi (**Figs 4G-4J**). No change in mRNA levels of the chemokine MCP-1 (**Fig 4L**) and of the anti-inflammatory cytokines, TFG-β and IL-10 were seen following ZIKV infection compared to mock animals (**S3A and S3B Figs**). To investigate whether ZIKV-induced inflammation of muscle tissue leads to an active process of atrophy, we evaluated the expression of Muscle RING Finger 1 (MuRF1) and Atrogin-1, both ubiquitin ligases which are associated to degradation of muscle proteins (23). We found an increased expression of both signaling proteins in the muscle of ZIKV-infected mice compared to the control (**Figs 4M and 4N**). These data support that ZIKV replication in muscle tissue promotes inflammation-induced muscular damage.

**Figure 4.**
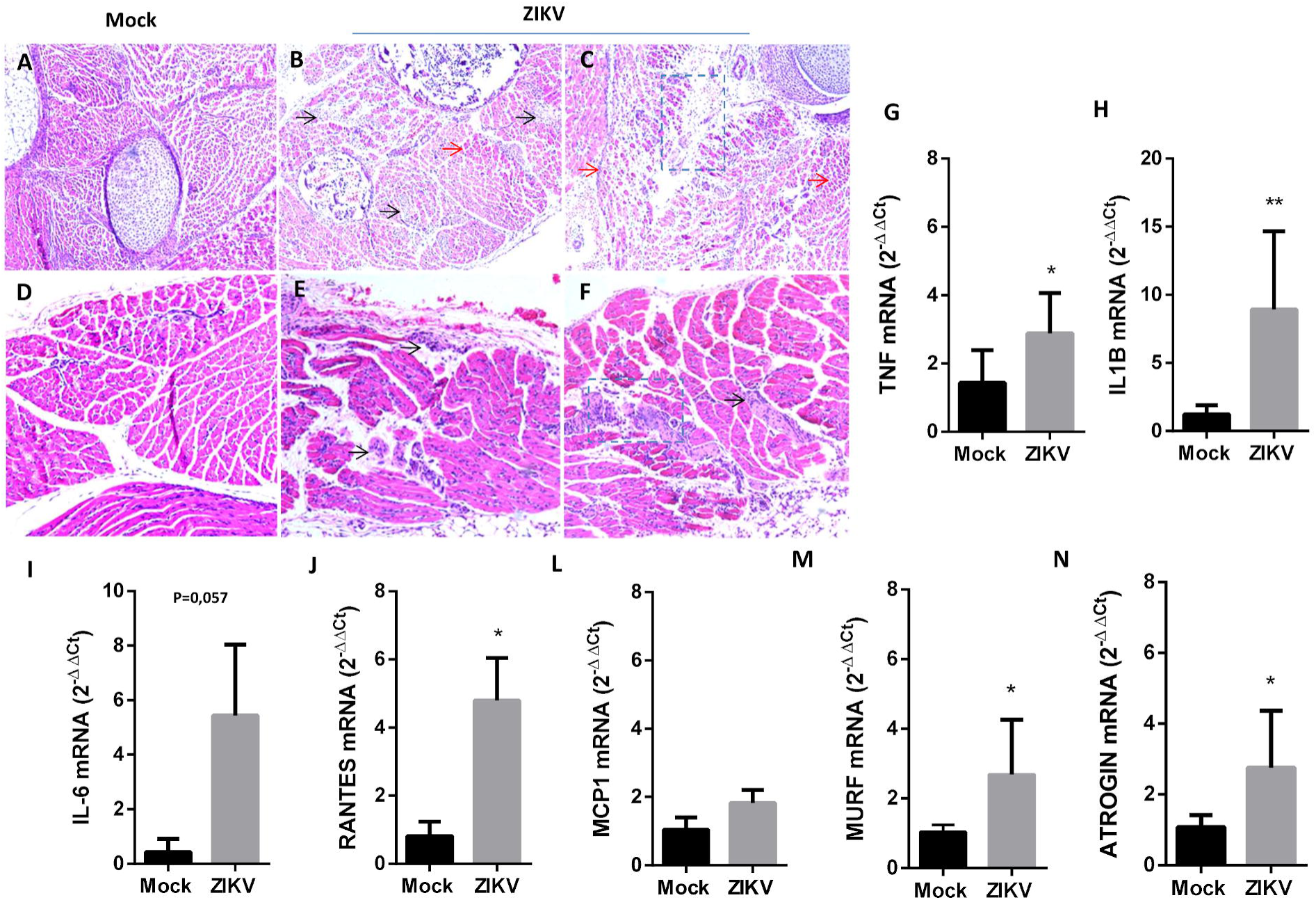
ZIKV induces muscle inflammation and damage in pups and neonate infection. SVA129 mice at 12.5 of gestation and 3 days-old WT SV129 mice were Mock or ZIKV infected. Skeletal muscle from hind legs of pups and of neonate were collected bilaterally and fixed at birth or 6 days post-infection, respectively, for histological analysis (**A-F**). Muscle tissues were embedded in paraffin after dehydration and tissue sections of 5 µm were prepared and stained with H&E. Scale bar = 100 μm. Black arrows areas of inflammation; red arrows indicate atrophy areas; dashed lines indicated areas of intense lesion. (**G-N**) Skeletal muscle from hind legs of neonates was collected at 6 days post-infection and the levels of TNF-α, IL-6, MCP-1, IL-1β, RANTES, MURF and Atrogin expression were determined related to Mock group by qPCR using β-actin expression as endogenous control. Values were plotted as mean ± Standard Error of Mean (SEM). Statistical analysis was performed using two-sided Mann-Whitney test.P *□0.05, **□0.01.

### ZIKV replicates in muscle prior to reach neural tissue in mice

To investigate the dynamics of ZIKV infection in mice tissues, we compared the time course of virus replication in neural and muscle tissues (**Fig 5A**). Viral amplification in muscle started immediately after virus inoculation (10-fold increase in viral titer is observed at 1 dpi), while an increase in ZIKV RNA levels in dorsal root ganglia (DRG), spinal cord (SC) and brain was detected only at day 4 of infection (ZIKV RNA levels were the same as the viral input until 2 dpi in these tissues, as indicated by dotted line in **Fig 5A**. The absence of virus replication in brain tissue at the early times after infection is consistent with the preservation of blood brain barrier (BBB) integrity in 3 days-old mice at the moment of ZIKV inoculation (**S3C Fig**), as well as after 2 days of infection (**S3D Fig)**. However, ZIKV amplification in the brain became evident at 4 days post-infection, but BBB integrity was not drastically disrupted at this time point (**S3E Fig)**. Despite the rapid amplification rates of ZIKV in muscle, the viral load in brain and SC reached levels higher than in muscle after 6 dpi, which was consistent with the known neurotropism of ZIKV.

**Figure 5.**
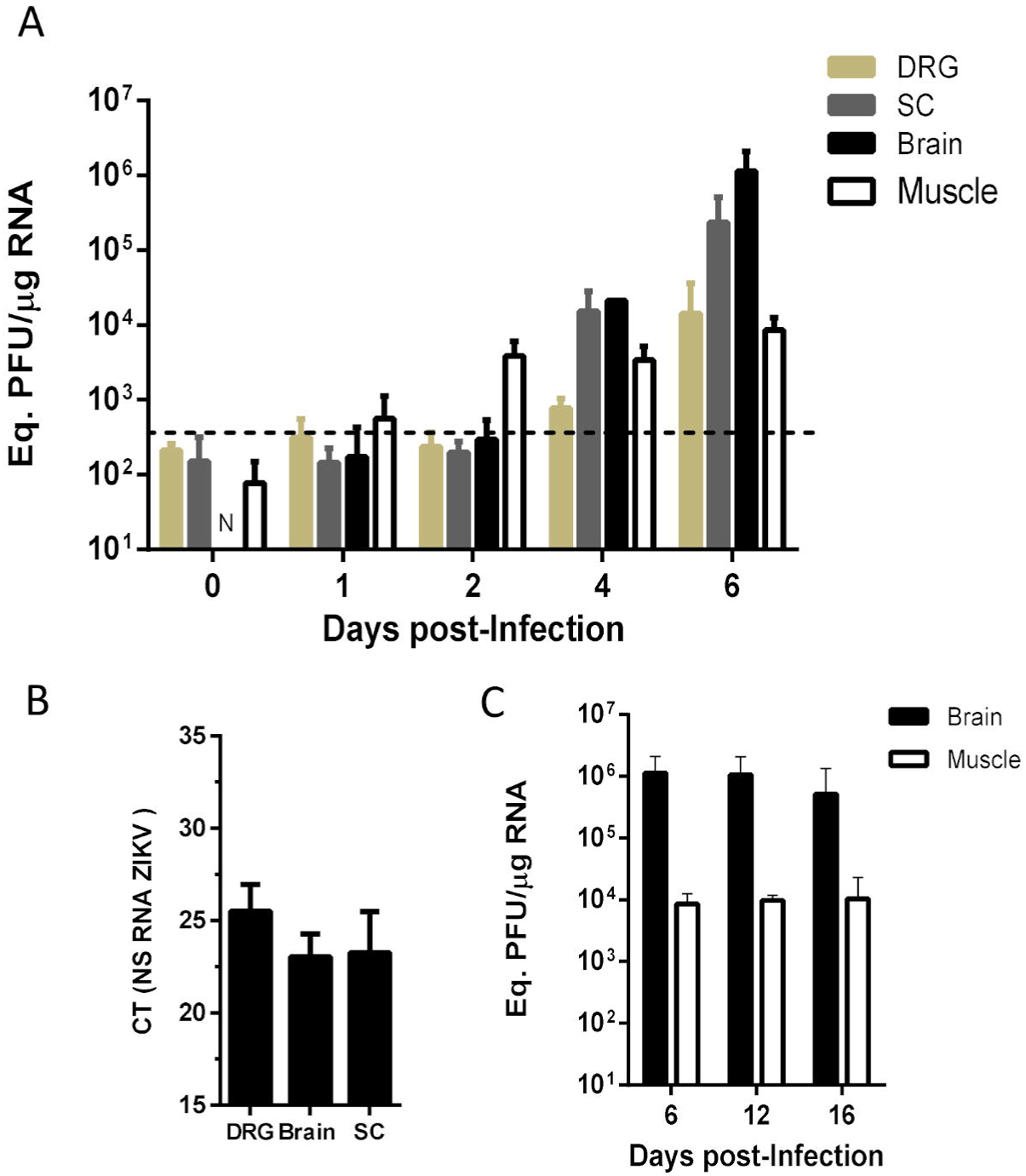
ZIKV replicates in muscle prior to reach neural tissue in mice. **(A**) 3 days-old WT SV129 mice were ZIKV infected and the ZIKV replication kinetic at muscle, brain, spinal cord (SC) and dorsal root ganglia (DRG) was determined by qPCR. (**B**) Negative strain amplification was detected by qPCR in neural structures 6 days post-infection and CT values (cycle threshold) were plotted. (**C**) ZIKV load was detected at late times post-infection in muscle and brain. ZIKV RNA load in **A** and **C** were plotted as mean ± standard error (SEM) of ZIKV equivalent of PFU/µg of total RNA.

Replication in DRG is detectable only after 6 days of infection. To confirm that the increase in ZIKV RNA in this tissue corresponds to an active viral replication, we searched for the negative strand of viral RNA (**Fig 5B**). Consistently with the data of pfu equivalent, lower levels of ZIKV RNA negative strand were detected in DRG compared to SC and brain. Interestingly, ZIKV could still be detected in muscle and brain tissue until 16 dpi at the same levels of 6 dpi in the last long-surviving mice (**Fig 5C**). The ZIKV titer was maintained with 2 logs of difference between these tissues. These data indicate that ZIKV establishes a rapid and sustained infection in the muscle tissue, which would contribute to viral amplification and dissemination to neural structures, the main target of replication.

## Discussion

ZIKV neurotropism has been extensively characterized due to its potential to induce microcephaly and other birth defects after gestational infection (24, 25). However, the contribution of ZIKV replication in peripheral tissues to pathogenesis is still unexplored. Thus, further investigations concerning the complete range of ZIKV induced lesions, as well as the determinants factors of pathogenesis severity are mandatory to prevent brain damage induced by the infection.

Muscular pain is a recurrent symptom associated with ZIKV infection, affecting up to 65% of patients in some outbreaks (26, 27). Skeletal muscle is a widespread body tissue comprised by mature fibers and myogenic precursor cells that are responsible for muscle development and repair (28). Muscle cells have been described as a target for replication of some arboviruses, such as Dengue (DENV) and Chikungunya (CHIKV), playing a significant role in their pathogenesis (29-31).Here we demonstrated that ZIKV replicates and damage cultured human primary muscle cells, and muscle tissue of fetus after maternal transmission and of neonate infected mice. Our data indicates that muscle is a target for ZIKV replication and site of injury. The ZIKV ability to infect and induce muscle damage is consistent with the high frequency of myalgia in patients, including those cases of self-limited disease and with neurological complications (32-34). In addition, it also suggests that an initial muscular replication may be a common step before ZIKV reaches neural structure.

Although ZIKV tropism in peripheral tissues was not well characterized yet, clinical studies have shown the viral persistence in several body fluids (21). First’s reports evaluating cell permissiveness demonstrated that ZIKV replicates at placental trophoblasts, endothelial cells, human skin fibroblast and neonatal keratinocytes (21, 35, 36). ZIKV also replicates and induce morphological alterations in human skin explants (35), that are consistent with skin lesions presented in many ZIKV-infected patients (37). After mosquito inoculation, skin cells (keratinocytes and dendritic cells) are the first target of ZIKV, followed by hematological dissemination. Mosquito or subcutaneous inoculation in non-human primates showed that ZIKV present a broad range tissue distribution with highest load at lymph nodes (38). Differing from DENV infection, *in vitro* and *in vivo* studies indicates that liver, despite susceptibility of some hepatocytes lineages (39), do not sustain high loads of ZIKV amplification in animal models (38). In agreement with this, we did not find ZIKV amplification in the liver during neonatal infection in mice. Our data suggest that the skeletal muscle is an important peripheral site for ZIKV amplification, mainly when compared with liver and spleen.

Fetal and neonatal muscle tissue is formed by fiber under maturation and muscle precursor cell, which differs according to its proliferative and fusogenic activity (40, 41). Similar to brain, were ZIKV replicates in neural progenitor’s cells (42, 43), we demonstrated that muscle precursor cell are the more susceptible to ZIKV than fibers, revealing that the preference for undifferentiated cell is a common feature of ZIKV tropism. In addition, ZIKV replication in neural stem cells also promotes cell death and decreases neurogenesis (44). Our findings suggest that ZIKV-induced cell death of muscle precursor cells may contributes to myogenesis disruption during fetal and neonatal muscle development. We demonstrated that ZIKV replication during muscle cells differentiation impact the number and area of the newly formed fiber. Impaired myogenesis could be related to cell death and with focal muscle atrophy observed in mice muscle at birth after fetal exposure to ZIKV. These data corroborate with clinical findings that showed atrophy of skeletal muscle in a human fetus after ZIKV maternal transmission (45). Further studies to evaluate the relationship between motor function and muscular replication of ZIKV should be address.

Muscle body composition after birth, despite present reduced myogenesis, grows very fast due to a high rate of protein synthesis (41, 42). However, a high number of precursor cells are maintained even at adult life, being responsible for muscle repair after injury (46). Thus, muscle precursor cells could be responsible by peripheral ZIKV amplification, even at adult life. In our neonate model, the load of ZIKV in muscle was sustained high until 16 days post-infection, raising the possibility of being a site of viral persistence and inflammation. ZIKV infection also promoted muscle lesions, with inflammatory cell infiltration, together with high levels of TNF, IL-1β, IL-6 and RANTES. Interestingly, the expressions of muscle degradation proteins were also increased during ZIKV infection in neonate mice. The increase in muscle mass loss could be trigger by several pro-inflammatory cytokines and others inflammatory mediator, such as TNF and IL-1β (47), that was fouded elevated in muscle during ZIKV infection, and could promote a reduction in muscle hypertrophy during development. In addition, the expression of anti-inflammatory cytokines that are associated with muscle tissue repair, such as TGF and IL-10 (48, 49), were not induced after lesion. Further analysis to evaluate long-term muscle injury and its function after ZIKV infection should be performed to explore this association.

Besides to be a peripheral site of ZIKV amplification, temporal analysis of muscular and neural ZIKV replication showed that the virus establishes a rapid and sustained kinetic of amplification in muscle tissue even before the viral replication in the neural tissues in mice. Furthermore, the inability of ZIKV disturb the BBB permeability, after early life infection, reinforces the possibility of viral accesses CNS by peripheral neurons. This is also supported by the detection of ZIKV negative strain in the DRG of neonates (6 dpi), and corroborates with previous studies that demonstrated ZIKV replication in explants and neurons from DRG of ZIKV-infected mice (16, 50). In addition, it was previously demonstrated that ZIKV reaches CNS without disrupt BBB by transcytosis, interfering on its permeability only at late stages of infection (15). However, it does not excludes that ZIKV could use peripheral neural route to access CNS, as already demonstrated to others neurotropic virus (20, 51).

Overall, our findings suggest that maintenance of viral load by muscle replication and inflammation could be an important step in ZIKV pathogenesis. It highlights the importance of investigating the molecular aspects associated with ZIKV-muscle interactions in the initial steps of the infection, dissemination and CNS damage in ZIKV-infected patients. Of note, the control of muscle viral replication could be an important issue for the development of therapeutic strategies to prevent neurological complications after ZIKV infection. Taken together our data contribute to a broader understanding of ZIKV pathogenesis and opens new aspects in the investigation of the long-term consequence of early infection.

## Material and Methods

### Virus propagation and quantification

Zika virus (ZIKV-BRPE, ref. KX197192), was propagated in C6/36 cell lineage cultured in L-15 medium (Leboviz’s Invitrogen) supplemented with 10% of fetal bovine serum (FBS), at 28°C. C6/36 cells were infected in a multiplicity of infection (MOI) of 0.01 in L-15 without serum for 1 hour. After adsorption, medium was removed and infected C6/36 was cultured for 7 days in L-15 supplemented with 5% of serum at 20°C. After this period, culture medium was collected and centrifuged at 2,000 x g for 10 min to remove cell debris. The clarified medium was aliquoted and stored at -80°C. The same procedure was performed in C6/36 uninfected cells, to production of Mock stock. Viral titer of the stock was determined by plaque assay in VERO cells.

VERO cells cultured in D-MEM high glucose (Invitrogen) without serum at 24-well plates at 37°C and 5% of CO_2_ atmosphere were infected with ten-fold serial dilutions of ZIKV stock for 1 hour. After this period, medium was replaced by the D-MEM high glucose medium with 1.5% of carboxy-methyl cellulose (CMC-Sigma Co.), 1% FBS and 1% penicillin / streptomycin (Invitrogen) and cells cultured for 5 days at 37°C and 5% of CO_2_ atmosphere. Cells were fixed with a solution of 10% Formaldehyde (Vetec, Sigma Co.) for 30 minutes, and then stained with crystal violet solution (20% Ethanol, 1% Crystal Violet (Sigma Co.) and H2O). After wash, viral stocks title was calculated as plaque forming units per mL (pfu/mL)

### Human primary muscle cells culture, differentiation and infection

HSMM cells (Human Skeletal Muscle Myoblasts) are precursor skeletal muscle cells isolated from the arm or leg of adult healthy donors, were obtained from Lonza cell facility (catalog number CC-2580). These cells were cultured following Lonza specification and medium (SkGMTM medium), at 37°C and CO_2_ atmosphere. For the induction of differentiation of myoblast into myotubes, cells were plated with high density and the SkGM medium was supplemented with 2% horse serum (GIBCO® - Life Technologies TM). The culture medium was replaced every two days until the formation of the fibers (about 5-6 days).

HSMM myoblast and myotubes were infected with ZIKV using a MOI of 5 in SkGM medium without serum during 1 hour. After infection, HSMM were cultured in SkGM medium supplemented with 5% of SFB. Culture medium was temporally collected to determine the amount of ZIKV released during myoblast and myotubes infection by plaque assay. Myoblat and myotubes viability was determined by 3-(4,5-Dimethylthiazol-2-yl)-2,5-Diphenyltetrazolium Bromide (MTT; Life Technologies) metabolization.

For the infection during differentiation HSMM was cultured as described above. After 1 day of differentiation stimuli (Day 1), HSMM were mock or ZIKV infected with MOI of 1 and cultured in SKGM differentiation medium until complete 5 days of stimuli (Day 5). Culture medium was temporally collected to determine ZIKV load at medium by plaque assay. Cells were fixed at day 1 and day 5 to immunostaining of fibers formed and cell viability was determined by MTT assay.

### Fluorescence microscopy

HSMM cells seeded in 24-well plates were fixed with 4% of formaldehyde in phosphate balanced salt solution (PBS - pH 7.4) at desired time post-infection. Staining for viral proteins was performed using conditioned media of 4G2 hybridome (anti-flavivirus mouse antibody) at 1:10 dilution. HSMM myofibers were stained using a polyclonal antibody against myosin heavy chain (MF20) at 1:50 dilution. After that, both were stained with a goat anti-mouse Alexa-fluor 488 (Invitrogen) at 1:500 dilution. Cells in each field were visualized with nucleus stained with DAPI (GIBCO® - Life Technologies TM) at 1:10,000 dilution. Images were acquired using an inverted fluorescence microscope (IX81 – Olympus) with magnification of 20X. Images of 4G2 positive cells of at least 10 fields were used to determine % of infected HSMM. The cell fusion index in cultured Mock and ZIKV-infected myotubes was determined by quantification of MF20 positive cells in % of total nucleus (DAPI) and fiber area was obtained measuring the fluoresce of MF20 stained normalized by cell number. Image quantification was performed using at least 10 fields using imageJ software (version 1.51 j.8).

### Mice infection and tissue sample

All experimental procedures performed were in accordance with protocol and standards established by the National Council for Control of Animal Experimentation (CONCEA, Brazil) and approved by the Institutional Animal Care and Use Committee (CEUA), from Federal University of Rio de Janeiro (protocol no. 014/16; CEUA-UFRJ, Rio de Janeiro, Brazil).

For mouse model of ZIKV maternal transmission we used 8 week-age type-I interferon receptor deficient (IFNAR-/-) pregnant mice in different gestational stages (**10.5, 12.5 and 14.5**). Pregnant mice were subcutaneously inoculated with mock or 10^5^ pfu of ZIKV in a final volume of 50 µL, and monitored until delivery. The pups were counted; vital signs were observed and photographed at post-natal day 0 (P0). Pups brain and skeletal muscle from left and right hind legs were collected at P0 and fixed in 4% formaldehyde to histological or stored at -80 °C to viral quantification analysis.

For neonatal mice model, three day-old wide-type (WT) SV129 mice were subcutaneously inoculated with mock or 10^6^pfu of ZIKV in the dorsum in a final volume of 50 µl. Each experimental group was housed individually with the uninfected mothering polypropylene cages, maintained at 25 °C with controlled humidity, under a 12 h light/dark cycle with free access to chow and water. Mice were monitored daily and body weight was measured every 2 days. Tissue samples of Mock- and ZIKV-injected mice were collected at different time points and stored at -80 °C until processed to qPCR analysis or fixed in 4% formaldehyde to histological analysis.

### Endothelial barrier integrity analysis

Neonatal mice were subcutaneously inoculated with 50 µL of 1 % Evans blue solution (VETEC) before (3 days-old, moment of infection) and 2, 4 and 12 days after the injection of mock or ZIKV. After 1 h, mice were perfused with PBS, and then the brain and other peripheral tissues (liver, spleen and muscle) were collected to observe Evans blue staining.

### Histology

Skeletal muscle from hind legs of mice were collected bilaterally at 6-days post-infection, fixed with 4% of formaldehyde and embedded in paraffin after dehydration. Paraffin-embedded tissue sections of 5 µm were prepared and stained with hematoxylin and eosin (H&E). Images were obtained using optical microscopy with a magnification of 10 X (Olympus BX40), and images were acquired using software Leica Application Suite 3.8 (Leica).

### Quantification of ZIKV RNA and proteins expression by qPCR

Tissues were homogenized using a fixed concentration (0.2 mg tissue/µl) in DMEM, and 200 µL of the homogenate were used for RNA extraction with Trizol (Invitrogen) according the manufacturer’s instructions. Purity and integrity of RNA were determined by the 260/280 and 260/230 nm absorbance ratios. One µg of isolated RNA was submitted to DNAse treatment (Ambion, Thermo Fisher Scientific Inc.) and then reverse-transcribed using the High-Capacity cDNA Reverse Transcription Kit (Thermo Fisher Scientific Inc).

For quantification of ZIKV RNA primers for ZIKV were used as described by Lanciotti (2008): forward, 5’-CCGCTGCCCAACACAAG-30; reverse, 5’-CCACTAACGTTCTTTTGCAGACAT-3’; probe, 5’-/56-FAM/AGCCTACCT/ZEN/TGACAAGCAATCAGACACTCAA/3IABkFQ/-3’ (Integrated DNA Technologies). Analyses were carried out on an Applied Biosystems 7500 RT–PCR system using the TaqMan Mix (ThermoFisher Scientific Inc) according to manufacturer’s instructions. Cycle threshold (Ct) values were used to calculate the equivalence of log_10_ PFU/µg of total RNA after conversion using a standard-curve with serial 10-fold dilutions of ZIKV stock.

To detect negative ZIKV RNA strand in dorsal root ganglion (DRG), RNA was extracted as described above and cDNA was synthesized using 2 pmol of ZIKV 835 forward primer instead of random primers. Real time quantitative PCR analysis was performed as described above.

Quantification of proteins expression at muscle was performed using the Power SYBR kit (Applied Biosystems; Foster City, CA). Actin was used as an endogenous control. Primer sequences were the following: IL-6 (FW-5’- TTCTTGGGACTGATGCTGGTG-3’ ; REV, 5-CAGAATTGCCATTGCACACTC-3’), MCP-1 (FW, 5’-GTCCCCAGCTCAAGGAGTAT-3’ ; REV, 5’- CCTACTTCTTCTCTGGGTTG-3’), RANTES (FW, 5’- GTGCCCACGTCAAGGAGTAT-3’ ; REV, 5’-CCTACTTCTTCTCTGGGTTG-3’), TNF (FW,5’-CCTCACACTCAGATCATCTTCTCA-3’ ; REV,5’- TGGTTGTCTTTGAGATCCATGC-3’), IL-1β (FW, 5’- GTAATGAAAGACGGCACACC-3’ ; REV, 5’-ATTAGAAACAGTCCAGCCCA -3’); Actin (FW, 5’-TGTGACGTTGACATCCGTAAA -3’; REV, 5’-GTACTTGCGCTCAGGAGGAG-3’). MURF1 (FW, 5’- GAGAACCTGGAGAAGCAGCTCAT -3’ ; REV, 5’-CCGCGGTTGGTCCAGTAG- 3’); ATROGIN-1 (FW, 5’-AGAAAAGCGGCACCTTCGT-3’ ; REV, 5’- CTTGGCTGCAACATCGTAGTT-3’).

### Statistical analyses

Statistical analyses were performed comparing means by two-tailed T-Student’s tests using Graph Pad Prism version 7.00 for Windows, Graph Pad Software, La Jolla California USA, www.graphpad.com.

## Acknowledgments

This work was supported by grants from Brazilian funding agencies: Fundação de Amparo à Pesquisa do Estado do Rio de Janeiro (FAPERJ); Conselho Nacional de Desenvolvimento Científico e Tecnológico (CNPq), Agência Financiadora de Inovação e Pesquisa (Finep) andCoordenação de Aperfeiçoamento de Pessoal de Nível Superior (CAPES), finance Code 001.

## Authors Contributions

IAM, DGL,CMF, JRC, CPF, CFB and LLdesigned the experiments. DGL, CMF, LGB, MOLS, SMBA, LDM and RLSN performed the experiments. IAM and DGL analyzed the results. IAM wrote the manuscript. JRC, CPF, CFB and ATDP reviewed and edited the manuscript.

## Supporting Information

**Supplementary figure 1.**
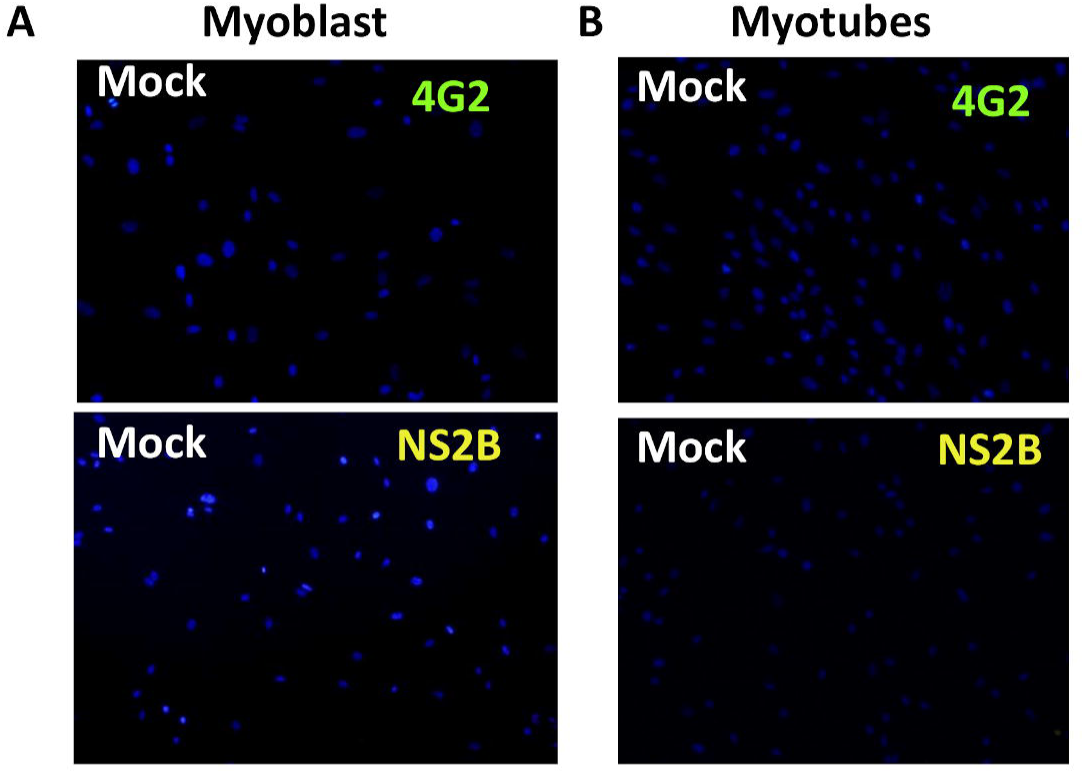
Immunofluorescence staining of ZIKV proteins in mock myoblast (A) and myotubes (**B**) 36 hours post-infection using anti-flavivirus E protein 4G2 and anti-NS2b of ZIKV.

**Supplementary figure 2.**
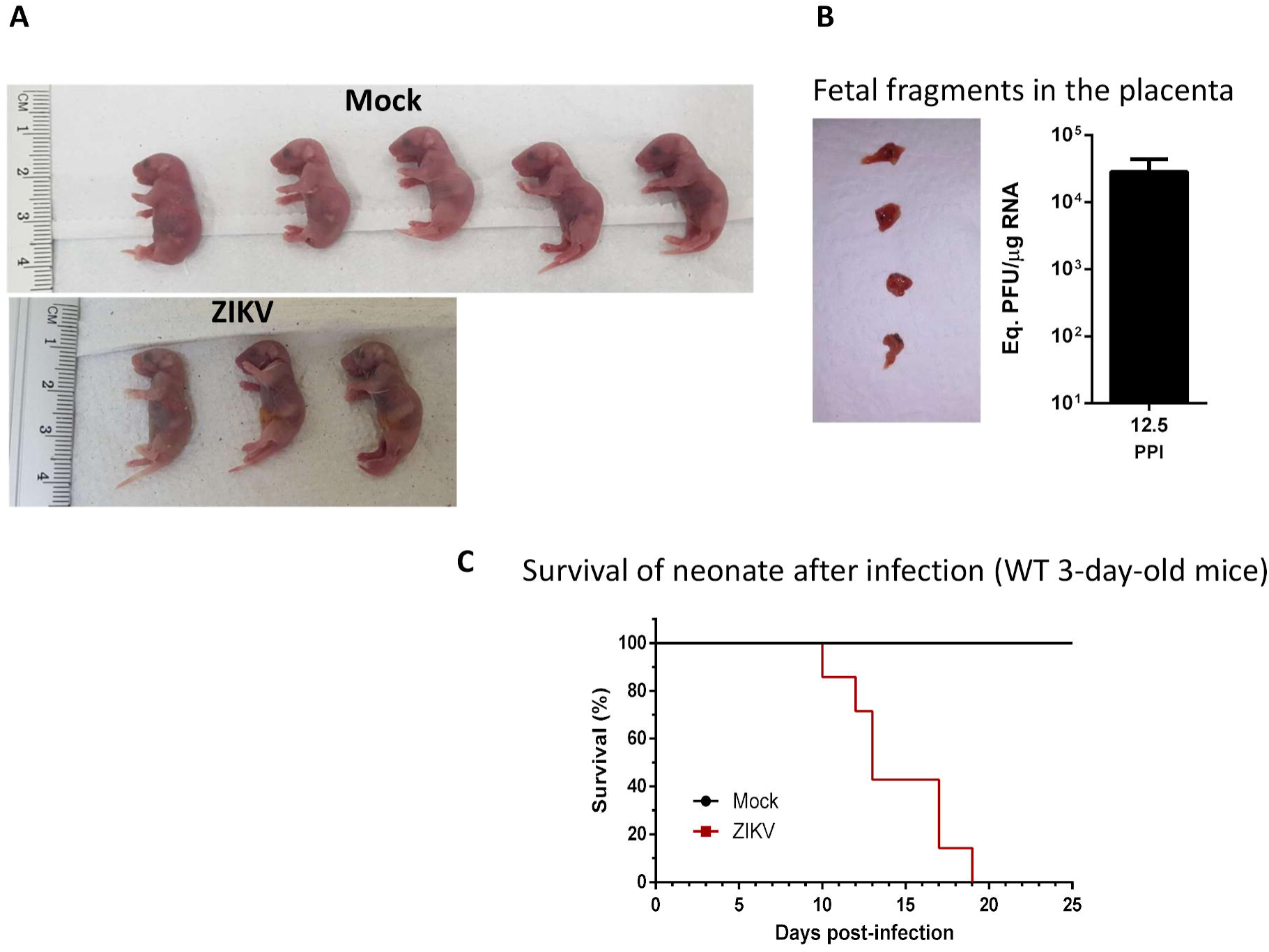
ZIKV infection during middle gestational stage in mice result fetal death. Females of IFNAR-/- mice were inoculated with ZIKV or mock on 12.5^th^ gestational day. (**A**) Representative images of litter at born from Mock and infected female. (**B**) Image of placenta of ZIKV infected female and for viral load at this samples were quantified by qPCR. (**C**) Survival curve of mock or ZIKV infected WT 3 days-old mice. The survival curve represents two experiments with 7 animals each.

**Supplementary figure 3.**
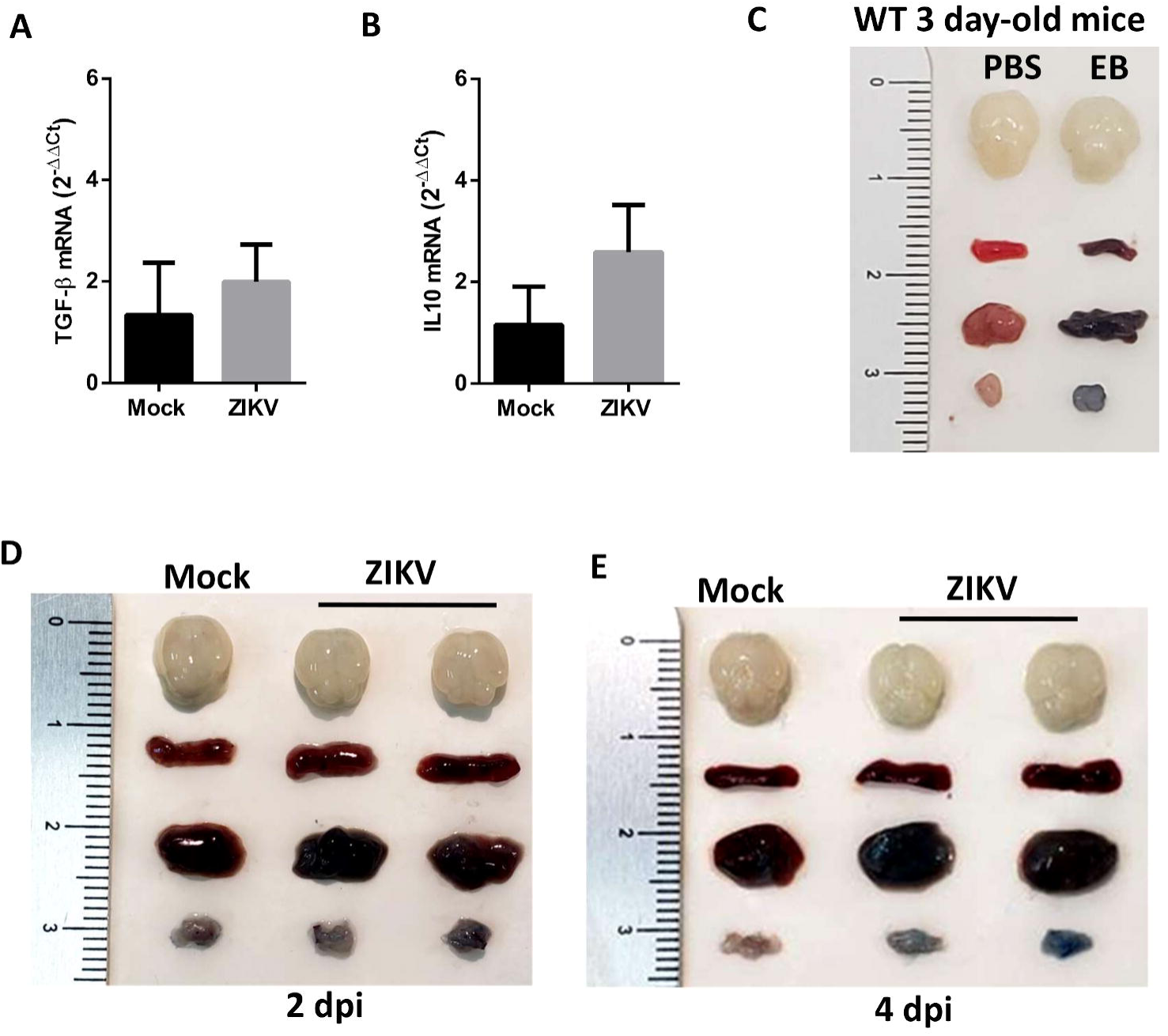
Skeletal muscle from hind legs of neonates was collected at 6 days post-infection and the levels of TGF-β (**A**) and IL-10 mRNA (**B**) were determined related to Mock group by qPCR using β-actin expression as endogenous control. Values are showed as mean ± Standard Error of Mean (SEM). Statistical analysis was performed using two-sided Mann-Whitney test. **B**lood-brain barrier permeability were analyzed in 3 days-old WT mice before infection (**B**), and at different times after Mock or ZIKV infection (**D-E**) was investigated with subcutaneously inoculation of 1% Evans blue solution (**EB**) or PBS. After a period of 1h animals were perfused with PBS, brains and peripheral tissues (spleen, liver and muscle) were removed for visualization.

